# Optimizing the Characterization and Quantification of Retinal Ganglion Cell Somas in Healthy and Injured Retinas Using Cellpose

**DOI:** 10.1101/2025.10.30.685421

**Authors:** Sarah E.R. Yablonski, Abigail Bishop, Sean D. Lydon, Ivan Zhu, Nicole Wang, Olivia J. Marola, Richard T. Libby

**Affiliations:** Department of Ophthalmology, Flaum Eye Institute, University of Rochester Medical Center, Rochester, NY, USA; Neuroscience Graduate Program, University of Rochester Medical Center, Rochester, NY, USA; The Center for Visual Sciences, University of Rochester, Rochester, NY, USA; Wisconsin IceCube Particle Astrophysics Center (WIPAC), University of Wisconsin-Madison, Madison, WI, USA; Department of Physics, University of Wisconsin-Madison, Madison, WI, USA; The Jackson Laboratory, Bar Harbor, ME, USA; Department of Biomedical Genetics, University of Rochester Medical Center, Rochester NY, USA

**Keywords:** Cellpose, glaucomatous neurodegeneration, optic nerve injury, automated quantification, automated characterization, retinal ganglion cell

## Abstract

Quantification of retinal ganglion cell (RGC) soma number and characterization of somal features are commonly used output metrics for investigation of optic neuropathies. Many investigators still perform these quantifications by hand, which is time consuming and prone to bias. Cellpose is an open-source Python package that can perform cellular segmentation and holds promise for automating somal analyses. Here, we designed a custom script incorporating the Cellpose package and custom-trained Cellpose models that are capable of automatic characterization and quantification of RGC somas. Our script, using our models, is capable of automatically counting RGC somas, along with characterizing RGC somal size. Further, we show that Cellpose can quantify RGCs using multiple cell-type specific markers and has the potential to quantify RGCs across the entire retina. Together, our custom Cellpose models and script which generates and analyzes Cellpose outputs provides a powerful tool for all-in-one RGC somal analysis.

## 1. Introduction

Retinal ganglion cells (RGCs) are the output neurons of the retina, with their projections supplying visual information from the eye to the brain. Death of RGCs is characteristic of optic neuropathies, such as glaucoma, which leads to visual loss and impairment of patients’ quality of life. In trying to understand the pathogenesis of these diseases, determine pathways underlying RGC death, and study potential therapeutics for preventing RGC loss, investigators commonly turn to counting and characterizing RGCs after various manipulations (e.g. ^1–3^). This process has evolved over time, but it is still common for investigators to complete these measurements by hand, which is time-consuming, prone to bias, and frequently limits the area of the retina that is quantified.

Processes such as quantifying RGC axonal density^4–7^ and interpreting optical coherence tomography data^8–10^ have had advances in the utilization of automated platforms to complete analyses. Similarly, RGC quantification has previously been performed automatically or semi-automatically^11–19^. These studies have done important work developing models that can quantify cells, identify cells using multiple markers (i.e. RBPMS, BRN3A), and perform quantitative analyses on soma size^13,14,20^. However, to the best of our knowledge, no system is yet able to combine all these features into one program, allowing for a comprehensive quantitative analysis of RGC somas. Cellpose is a software package for automated cellular segmentation that allows users to train models tailored to their own data^21,22^. Many distinct fields, including neuroscience, have utilized this software for segmentation and quantification of relevant features (e.g. ^23–26^). Here, we created a custom script utilizing the Cellpose platform to rapidly identify, quantify, and characterize RGCs in naïve states and in instances of RGC death.

## 2. Materials and Methods

### 2.1 Mice

C57BL/6J (B6), DBA/2J (D2), and D2.*Ddit3^-/-^*^27^ mice were used; B6 and DBA/2J mice were originally purchased from The Jackson Laboratory (Stock# 00664 and 000671, respectively) and were bred and maintained in-house. All mice included in the study were older than 1.5 months. Both male and female mice were used. Mice were fed chow and water *ad libitum* and housed on a 12-hour light-to-dark cycle. All experiments were conducted in adherence with the Association for Research in Vision and Ophthalmology’s statement on the use of animals in ophthalmic and vision research and were approved by the University of Rochester’s Committee on Animal Resources. Some of the images used throughout the publication have been previously published for use in investigating biological questions not addressed in this manuscript^27^. Images from DBA/2J and D2.*Ddit3^-/-^*mice, along with the mouse handling and immunofluorescence procedures were outlined in a publication outlining the role of *Ddit3* in mouse glaucomatous neurodegeneration models^27^. These methods will not be addressed in this manuscript.

### 2.2 Controlled Optic Nerve Crush (CONC)

Controlled optic nerve crush (CONC) was performed as previously described^28–30^. Briefly, mice were anesthetized using 100 mg/kg ketamine and 10 mg/kg xylazine delivered intraperitoneally. The analgesic meloxicam (2 mg/kg) was delivered subcutaneously prior to surgery. The optic nerve was exposed and crushed immediately behind the globe for five seconds with self-closing forceps. The contralateral eye received a sham surgery, where the optic nerve was exposed but not crushed. Antibiotic ointment was applied to the eye following the procedure. Eyes were harvested 5 days or 14 days following CONC.

### 2.3 Immunofluorescence

As previously described^27,29,30^, eyes were harvested and fixed in 4% PFA in 1X PBS for 90 minutes. Retinas were dissected out of the optic cup and blocked in 10% horse serum with 0.3% Triton^TM^ X-100 (Fisher scientific, 9002-93-1) in 1X PBS overnight at 4°C. Primary antibody solution again containing 10% horse serum with 0.3% Triton^TM^ X-100 was then applied to the retinas for three days at 4°C. Primary antibodies included rabbit anti-RBPMS (Genetex, GTX118619, 1:250), mouse anti-Brn3a (Santa Cruz Biotechnology, 14A6, 1:200) and mouse anti-Tubulin β3 (TUJ1) (BioLegend, 801201, 1:1000). Following washes with 1X PBS, retinas were then incubated for two days at 4°C in a secondary antibody solution diluted into 1X PBS. Secondary antibodies included Alexa Fluor™ donkey anti-rabbit 647 (Invitrogen, A21447, 1:1000) and Alexa Fluor™ donkey anti-mouse 555 (Invitrogen, A31570, 1:1000). Retinas were then washed, cut into four quadrants, and mounted on microscope slides with the ganglion cell layer facing upwards. Retinas were mounted using Fluorogel in TRIS buffer (Electron Microscopy Sciences, 17985-11).

### 2.4 Fluorescent Imaging and Manual Cell Quantification

Individuals taking images and quantifying images were masked to experimental group. Images were taken using a Zeiss Imager M1 fluorescent microscope at 40X magnification. Eight images per retina (two per quadrant) were taken approximately 500 μm from the peripheral edge of the retina, as previously described^31–34^. Manual quantification of RBPMS+ and TUJ1+ cells was performed using the cell-counter tool in ImageJ. Four trained evaluators counted RBPMS+ images, and three trained evaluators counted TUJ1+ images.

Soma size of RBPMS+ cells was completed in ImageJ by thresholding the image and converting images to binary. The watershed function was used to create boundaries between cells and the analyze particles function was used to obtain the final soma size measurement. Soma size measurements were independently completed by two trained evaluators. Cellular counts and soma size analyses were timed by the individuals completing them.

Images of entire retinas (Figure 6) were taken using a Keyence BZ-X800 epifluorescent microscope at 10X magnification. Images were stitched together using associated Keyence analysis software.

### 2.5 Cellpose Model Development and Testing

Python version 3.8.5 or later and Cellpose version 2.2 were used. Automatic detection of RBPMS+ and TUJ1+ cells was completed using custom-trained models designed for the Cellpose algorithm^21,22^. Cellpose models were trained using the human-in-the-loop approach outlined by Pachitariu and Stringer^22^. The “CP” model provided in Cellpose version 2.2 was used as the base model and all other modeling parameters were unchanged during development. 32 images from four eyes were used to train both the RBPMS+ and TUJ1+ models, with eyes from all relevant experimental groups being represented in the data set (Figure 1). 8 images from one retina were trained for the BRN3A Cellpose model. A small region of a full-retina image (approximately 2 mm by 200 μm and spanning from the periphery to the central retina) was used to train the full-retina RBPMS Cellpose model. Further, we developed a custom Python script that simultaneously calculates multiple RGC outcome metrics using Cellpose outputs. Our script was developed using Python, Cellpose and all packages used to run Cellpose. OpenCV2 incorporated into the script to calculate the length and width of each cell identified by Cellpose.

**Figure 1:**
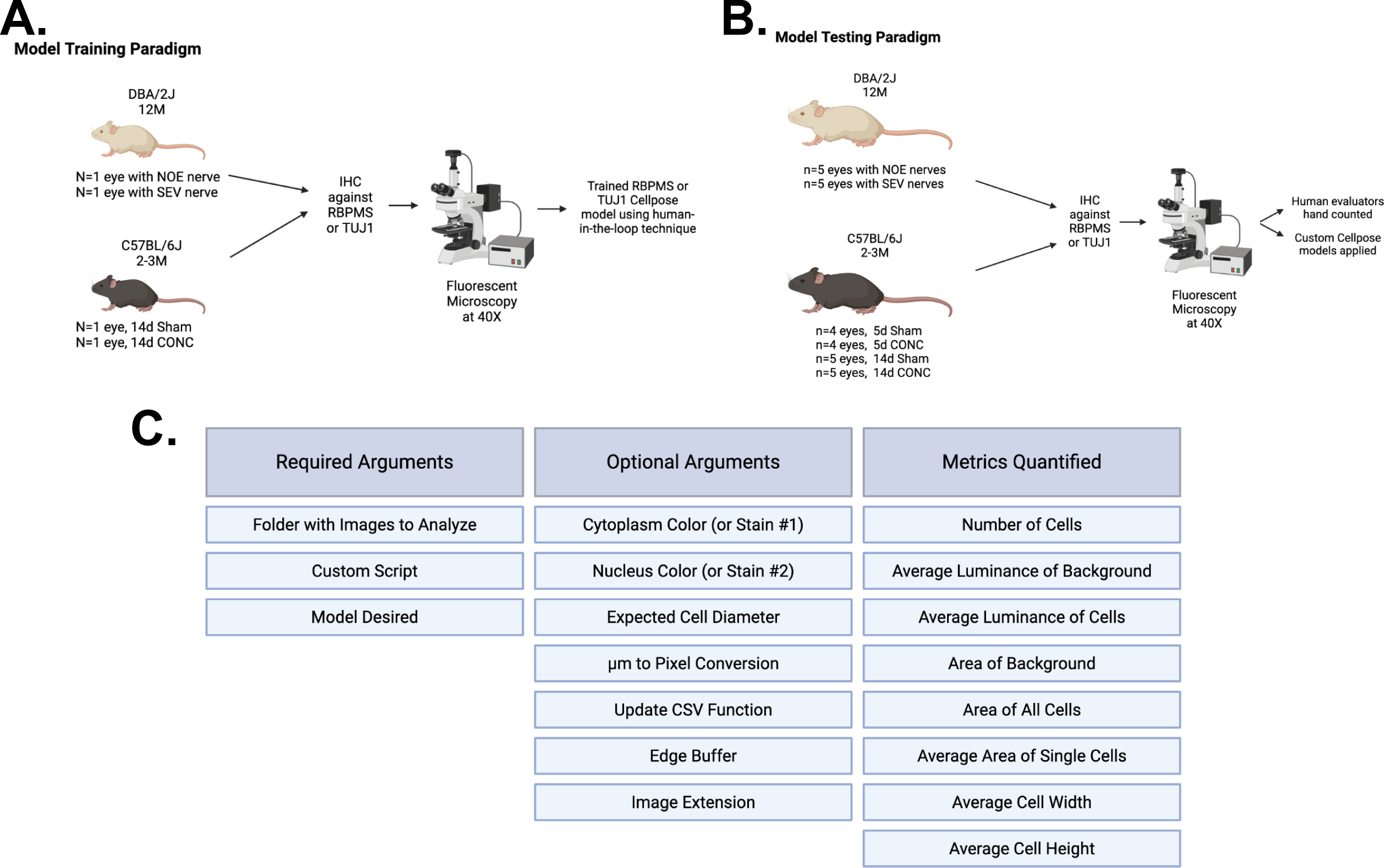
Developing custom Cellpose models and scripts for quantifying RGCs. **A.** Schematic outlining how custom TUJ1 and RBPMS models were developed within the Cellpose platform. **B.** Schematic outlining how custom TUJ1 and RBPMS Cellpose models were tested. **C.** Table outlining the features of the custom script written to use with Cellpose. Required arguments for script function, optional arguments to adjust data analyzed, and all metrics quantified and displayed in the .csv file are shown. BioRender was used to create schematics.

A separate set of 224 images was used to validate the RBPMS and TUJ1 models. This set of images was first analyzed by trained human evaluators and then processed with the Python script using the custom-designed Cellpose models for comparison. Analysis completion time was recorded by both the script and by each of the trained evaluators. The images were processed in Cellpose on multiple computers including a MacBook Pro 2020 with a 1.4 GHz Quad-Core Intel® Core™ i5 processor and Intel Iris Plus graphics, a Dell Precision 7540 with Intel® Core™ i9 Processor and NVIDIA Quadro RTX3000 Graphics, and a Windows Desktop computer with a 2.9 GHz Intel® Core™ i5-10400 Processor and NVIDIA GeForce GTX 1050 Ti graphics. To validate the BRN3A Cellpose model and the full-retina RBPMS model, separate images and retinas were used from the original training images. Note, although all models and data presented here were designed using Cellpose 2.0, Cellpose 3.0 was released during preparation of the manuscript^35^. Our script is compatible with Cellpose versions 2.2 through 3.1.1.2, as are our Cellpose models.

### 2.6 Statistical Analysis

Data were analyzed using GraphPad Prism 10 and Matplotlib^36^ software. Data from trials comparing Cellpose to the average human evaluator were analyzed using a simple linear regression. Data from trials comparing multiple groups (e.g. Cellpose vs Human Evaluators in specific experimental conditions, Figs. 3&4) were analyzed using a two-way ANOVA followed by a Fisher’s LSD test. Intraclass Correlation Coefficient (ICC) between human evaluators was determined by two individuals using formula ICC(A,1), as defined by McGraw and Wong^37^. *P* values < 0.05 were considered statistically significant and data is reported as mean ± standard error of the mean (SEM) throughout the manuscript.

## 3. Results

### 3.1 Creating Custom Cellpose Models and Script to Quantify RGCs

To analyze RGCs using the Cellpose platform effectively, custom Cellpose models and a custom script were developed (Figure 1). Using custom Cellpose models allowed for the analysis of RGCs using two RGC markers, RBPMS and TUJ1, which were chosen based on their relevance to previous literature characterizing RGC somas (e.g. ^1,2,38–40^). Additionally, data from both healthy and injured retinas were incorporated into Cellpose model training to ensure there was not a bias toward health status of RGCs during quantification (Figure 1A). Injured retinas included those from DBA/2J mice with chronic ocular hypertension and those from controlled optic nerve crush (CONC). These injuries were chosen because they are common systems used to study both RGC loss and neuroprotection (e.g. ^27,31,32,41–49^). DBA/2J retinas used in both the training and testing of Cellpose models were from mice aged to 12 months and whose corresponding optic nerves had previously been judged to have no histologically detectable damage (NOE, as previously defined^50^) or serve (SEV) degeneration (greater than 50% axonal loss^50^). Some data from these retinas were previously published^27^. An injured and a contralateral sham retina, obtained 14 days post-CONC, were also included in the model development image set. 32 images were used in total, and once the images were compiled, the Cellpose models were then trained using a human-in-the-loop methodology^22^ (Figure 1A).

To test whether the Cellpose models were comparable to human evaluators, a separate dataset of images was analyzed. Animal models included in this dataset were 12-month DBA/2J, 5-day CONC or sham, and 14-day CONC or sham (Figure 1B). 224 total images were included in the test dataset. Each image was first manually evaluated by four trained individuals counting the RBPMS dataset and three trained individuals counting the TUJ1 dataset. Following the manual counts, the number of cells in all images was then counted by the using Cellpose and the appropriate custom-trained Cellpose model for comparison.

Along with the custom trained models, a custom script was developed that allowed for easy data processing and additional analyses to be completed (Figure 1C). This script can be run directly from the command line and only requires the user to specify a folder of images to analyze and a model to analyze the images with. Optional arguments can be used to customize the analysis process to a given need or image set. The output of this script includes both the Cellpose segmentation file for each image (ending in .npy) and a .csv file containing all the data calculated by Cellpose for each image (Figure S1). By default, the script computes multiple supplemental RGC parameters for each image, without the need for additional analyses. These parameters include the average area of individual cells (or soma size), the average length & width of individual cells, and the average luminance of cells.

### 3.2 Applying Custom Cellpose RBPMS Model to Quantify RGCs

The 40X RBPMS Cellpose model reliably identified RBPMS+ RGCs (Figure 2A). Importantly, RBPMS is known to stain RGC somas and, more weakly, endothelial cells withing the ganglion cell layer^1^. The RBPMS Cellpose model was also able to distinguish between these two cell types and only counted the RGCs (see figure 3F, for example). Additionally, there was a high correlation between the human evaluator’s average cell count per image and the custom script’s (using the RBPMS Cellpose model) count across the entire dataset (Figure 2B, R^2^ =0.9961), validating that the script could quantify RBPMS+ RGCs as well as a human evaluator. Among the human evaluators, there was variation in how each individual image was counted (Table 1, ICC). To understand the scope of this variation and whether the counts obtained using the Cellpose model were within the human variation, a percent difference plot was made (Figure 2C). In this plot, data points for each image were ordered by increasing cell count (reduced damage), as calculated by the human evaluators. The variability of the human evaluator counts is shown by via the grey band. For each image, the grey band is bounded by the maximum and minimum number of cells counted by a human evaluator, expressed as the percent difference relative to the average of all human evaluator counts for the same image. Across most images, the variation in counts calculated by human evaluators remained relatively low. The script’s count for each image, also expressed as a percent difference relative to the average count from human evaluators for that image, is overlayed as a black dot. This revealed that while the counts obtained using the Cellpose and the custom RBPMS model frequently fell within the human variation for a given image, there were also occasional examples of Cellpose over- or under-counting the number of cells in a given image compared to the ave3rage human count (Figure 2C).

**Figure 2:**
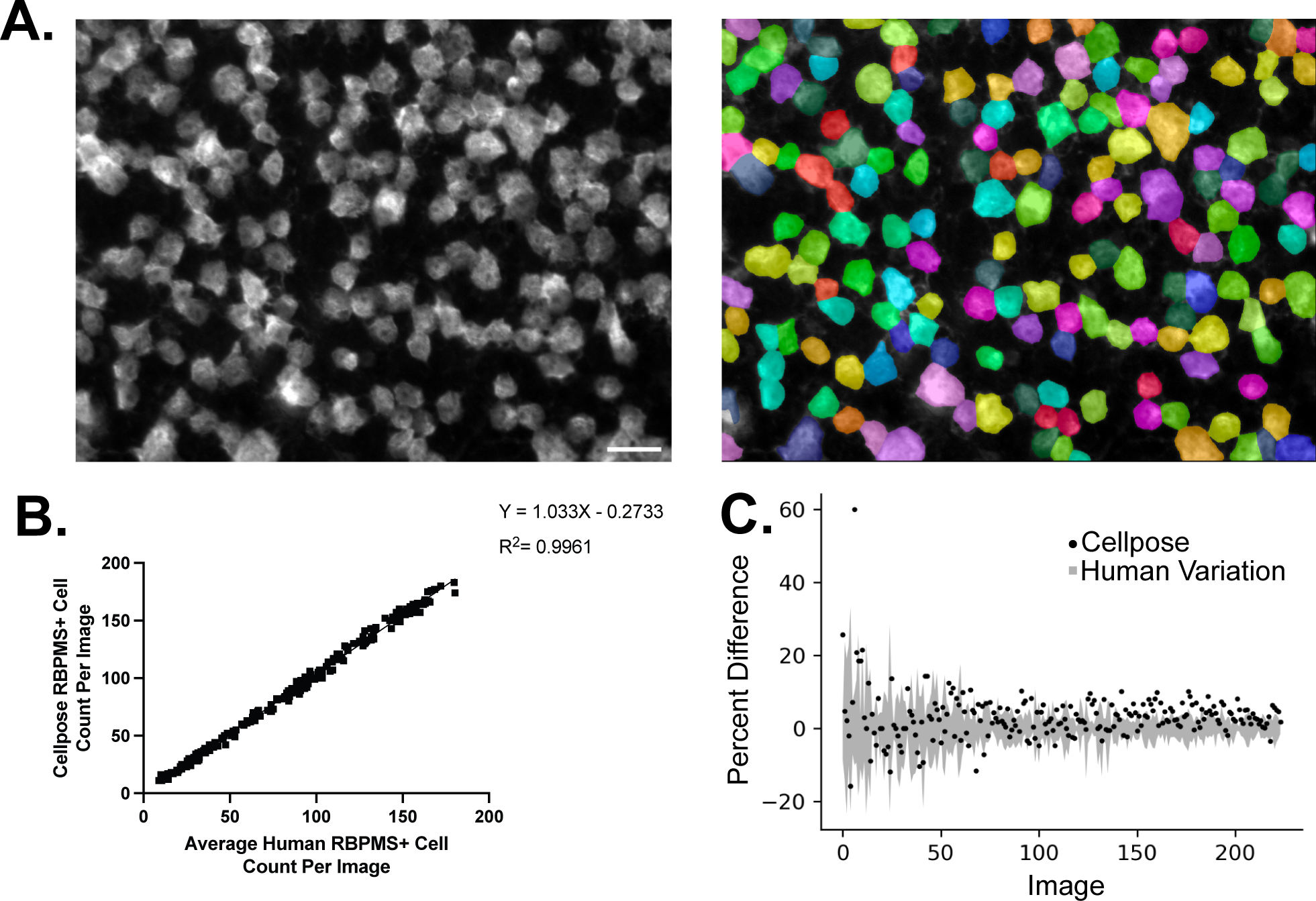
RBPMS Cellpose model has high correlation with human evaluators. **A.** Example RBPMS-labeled RGCs shown in their original image (left) and with superimposed Cellpose-identified cellular boundaries (right) after running the custom-trained RBPMS Cellpose model. Scale bar = 20 μm. **B.** Simple linear regression analysis comparing the custom script’s (running the RBPMS Cellpose model) quantifications of RGCs to the average human evaluator’s quantifications of the same images. There is a high degree of correlation. Four trained human evaluators assessed each image, and 224 images were quantified. **C.** The minimum and maximum human count for each image was compared to the average human count for each image (gray bars), represented as percent difference. The percent difference between the script’s counts and the average human evaluator’s count is also shown (black dots). Images are ordered from lowest count to highest count (most damage to least damage).

**Figure 3:**
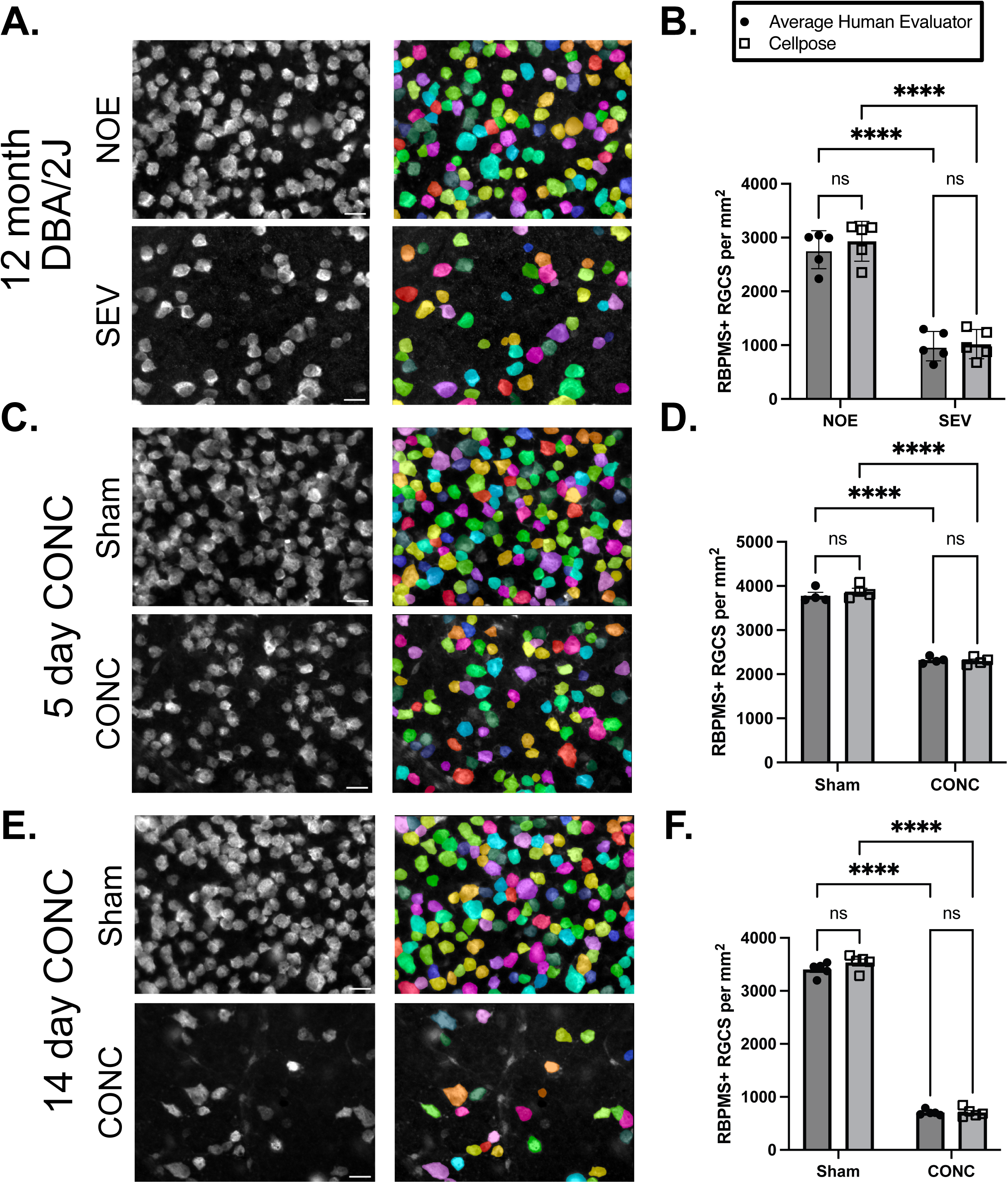
The script accurately quantifies RBPMS+ RGCs in different glaucoma-relevant models. **A.** Retinal flat mounts from eyes corresponding to optic nerves with NOE or SEV degeneration from 12-month DBA/2J animals. Flat mounts were labeled for RBPMS (left). Cellpose-identified cellular boundaries were superimposed onto images using custom trained RBPMS model (right). **B.** The script accurately quantifies RGC density in NOE and SEV groups in a manner comparable to human evaluators (P<0.001 for both significant comparisons, n=5 eyes per group, n=4 human evaluators assessed each image, two-way ANOVA, Fisher’s LSD post-hoc). **C.** Retinal flat mounts labeled for RBPMS 5-days post CONC (left). Cellpose-identified cellular boundaries were superimposed onto images using custom trained RBPMS model (right). **D.** The script accurately quantifies RGC density in sham and CONC groups in a manner comparable to human evaluators (P<0.001 for both significant comparisons, n=4 eyes per group, n=4 human evaluators assessed each image, two-way ANOVA, Fisher’s LSD post-hoc). **E.** Retinal flat mounts labeled for RBPMS 14-days post CONC (left). Cellpose-identified cellular boundaries were superimposed onto images using custom trained RBPMS model (right). **F.** The script accurately quantifies RGC density in sham and CONC groups in a manner comparable to human evaluators (P<0.001 for both significant comparisons, n=5 eyes per group, n=4 human evaluators assessed each image, two-way ANOVA, Fisher’s LSD post-hoc). Scale bars, 20 μm.

**Table 1.** Additional statistics comparing the average human evaluator to Cellpose for the RBPMS dataset. Numbers shown represent average ± SEM.

|  | Minimum Count | 25% Percentile of Counts | Median Count | 75% Percentile of Counts | Maximum Count | Average Time to Analyze Dataset (minutes) | ICC between Human Evaluators |
| --- | --- | --- | --- | --- | --- | --- | --- |
| Average Human Evaluator | 8.8 | 32.3 | 86.4 | 128 | 180.3 | 242.7±38.9 | 0.995 |
| Cellpose | 11 | 34.3 | 88.5 | 132 | 183 | 48.7±7.3 |  |

Given that the script counted RBPMS+ RGCs in a manner comparable to human evaluators, we next tested its performance on two different retinal injury models: DBA/2J and CONC (both 5- and 14-days post injury). 12-month-old DBA/2J mice typically have optic nerve degeneration and RGC somal loss^28,48^. However, this damage is variable, and some optic nerves show no histological signs of glaucomatous neurodegeneration (NOE), whereas others show severs (SEV) degeneration^48^. When retinas corresponding to optic nerves showing either NOE or SEV degeneration were tested using the RBPMS Cellpose model, Cellpose was able to identify RBPMS+ RGCs in both conditions (Figure 3A). This was also represented in the RGC density calculations, which showed that the script did not differ from human evaluators in either the NOE or SEV groups (Figure 3B).

The peak of RGC death occurs at 5 days following CONC^29^, making this an ideal timepoint to study processes involved in RGC somal death^51,52^. Similar to DBA/2J retinas, the RBPMS Cellpose model was able to accurately identify RBPMS+ RGCs in both the sham and CONC groups (Figure 3C). Again, the RGC density did not differ between the script and human evaluators for the sham group or the CONC group (Figure 3D). Additionally, the script was able to identify the same difference in RGC density as the human evaluators between the sham and CONC groups (Figure 3D). Finally, we used the script to count RGCs 14 days after CONC. At this timepoint, the majority of RGC death has completed, making this an important timepoint for many researchers studying whether specific genetic manipulations or treatments improve RGC survival^27,47,51,53,54^. As before, the Cellpose RBPMS model was successfully used to identify RBPMS+ RGCs in both the sham and CONC groups (Figure 3E). The sham and CONC RGC density did not differ from those of human evaluators, and the script was able to identify the difference in RGC density between the sham and CONC groups (Figure 3F). Of note, images included in the test dataset represent the mid-periphery region of the retina. When Cellpose and the RBPMS model was tested on RBPMS+ RGCs present in different regions of the retina, high levels of performance continued to be achieved (Figure S2). Together, these data show that the RBPMS Cellpose model combined with the custom script can accurately identify and quantify RGCs in a variety of different injury states.

Further, another important goal of developing the custom script was to ensure it would save investigators time during the data analysis pipeline. The three quickest human evaluators counted the RBPMS test data set in 188, 222, and 318 minutes (for an average time of 217.6 minutes). In comparison, running the entire RBPMS dataset through the script took three separate machines with different processing power (see methods for details) 39, 44, and 63 minutes (average time 48.7 mins; Table 1). Further, while the script is running, investigators do not need to be present at their computer, meaning the only time needed to complete the analyses is the time needed to start the program. This represents a substantial amount of time saved by analyzing the data via the script instead of by hand. Additionally, this single run of the script also provided multiple additional RGC output metrics, including mean soma size in each image (see below). These metrics would otherwise have to be determined by a time intensive, manual processing. Soma size calculations took an average of 84.2 additional minutes for human evaluators to complete, greater than the entire time needed to run the dataset through the script. Taken together, the script combined with the RBPMS Cellpose model is comparable to the average human analyzer, can analyze RGCs in different injury states, and saves investigators time when completing their somal analyses.

### 3.3 Applying Custom Cellpose Models to Quantify RGCs Labeled for Additional RGC markers

RBPMS is a popular marker to characterize RGC somas, but other labels are also widely used. Another antibody commonly used in RGC quantification studies is TUJ1^2^. Unlike RBPMS which is specific to RGC somas, TUJ1 labels RGC somas, axons, and dendrites. To test whether Cellpose could be used to quantify RGCs labeled by TUJ1, a custom Cellpose TUJ1 model was created and tested. As with the RBPMS Cellpose model, the TUJ1 Cellpose model was created and trained using 32 images from both healthy and injured retinas, then tested using 224 images from every experimental group represented in the study (Figure 1A-B). Sample output shows reasonable identification of RGCs using the Cellpose TUJ1 model (Figure 4A). However, Cellpose’s TUJ1 RGC count was not as closely correlated to counts from human evaluators as Cellpose’s RGC counts with the RBPMS model (Figure 4B, R^2^=0.8848). Additionally, there was a wider variation between the human counts for each TUJ1+ image (Figure 4C). Despite this wider variability between human counters, the script’s counts commonly fell outside of the range of human variation, especially in images with a higher number of RGCs (Figure 4C). This imprecise evaluation of TUJ1+ images with high RGC density was demonstrated clearly in the 14-day CONC dataset. Cellpose’s RGC density estimate was similar to the human average in the CONC group, but significantly lower in the sham group (Figures 4D-E). Although the difference in RGC density between sham and CONC groups was still significant when assed by Cellpose, the undercounting of TUJ1+ RGCs in the sham group suggests that using this custom Cellpose TUJ1 model would result in an inaccurate representation of the dataset. Cellpose results had similar inaccuracies when counting TUJ1+ cells 5-days after CONC (Figure S3). Cellpose showed comparable RGC density estimates to human analyzers for the 5-day CONC group, but the Cellpose counts for the sham group were significantly lower than the human average. Upon review, the TUJ1 Cellpose model missed multiple cells that the average human could identify correctly, likely explaining the difference. Interestingly, the Cellpose script was able to accurately identify and quantify TUJ1+ RGCs in the 12-month DBA/2J dataset in eyes corresponding to both NOE and SEV nerves (Figure S3). Taken together, using the TUJ1 Cellpose model with Cellpose may accurately quantify RGCs at low RGC densities, but the model loses accuracy as RGC density increases.

**Figure 4:**
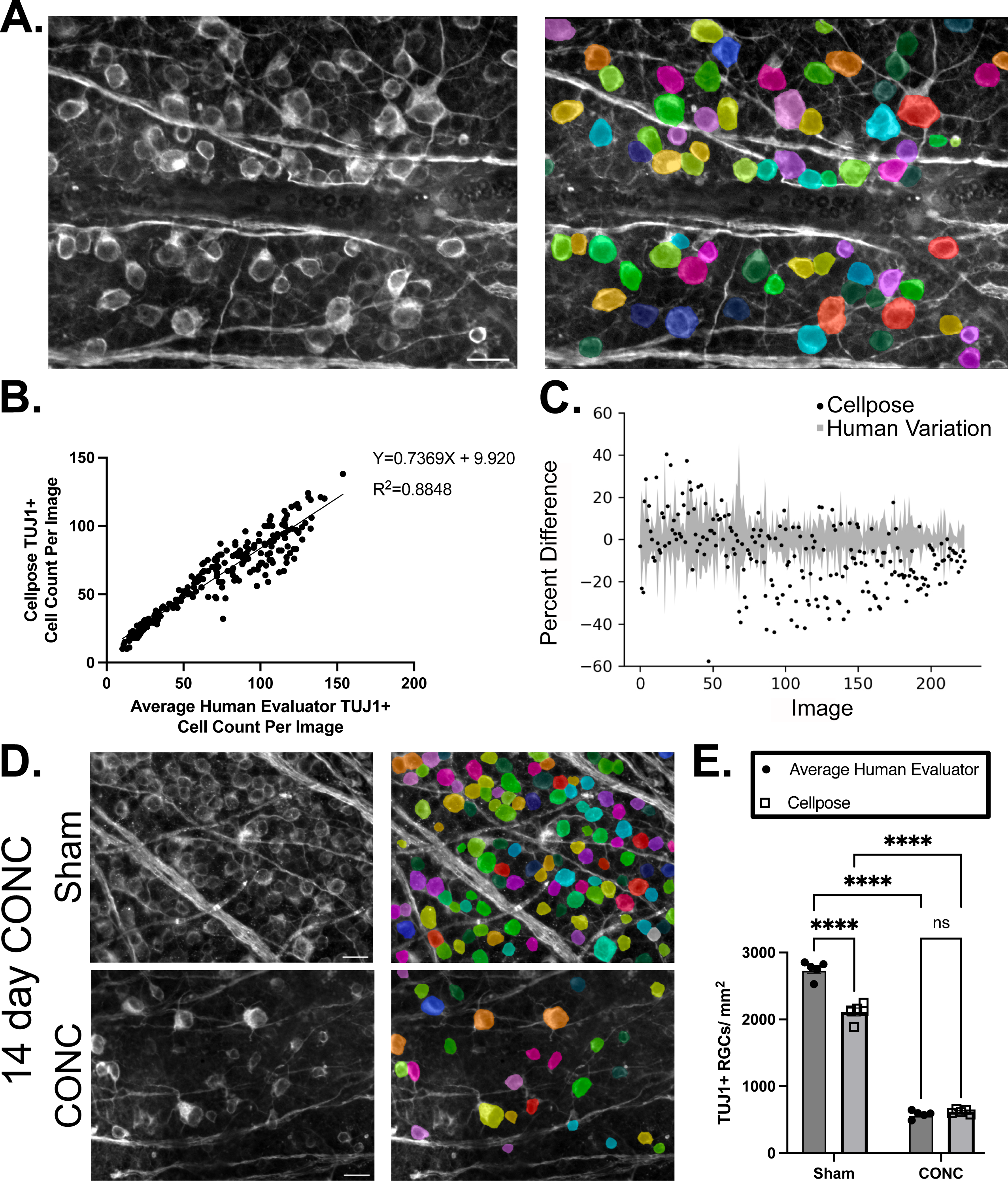
TUJ1 Cellpose model is less comparable to human evaluators in assessing RGCs. **A.** Example TUJ1-labeled RGCs shown in their original image (left) and with superimposed Cellpose-identified cellular boundaries (right) after running the custom-trained TUJ1 Cellpose model. **B.** Simple linear regression analysis comparing Cellpose’s quantifications of TUJ1+ RGCs to the average human evaluator’s quantifications of the same images. A moderate correlation between the two groups is demonstrated. Three trained human evaluators assessed each image, and 224 images were quantified. **C.** The minimum and maximum human count for each image was compared to the average human count for each image (gray bars), represented as percent difference. The percent difference between the Cellpose’s counts and the average human evaluator’s count is also shown (black dots). Images are ordered from lowest count to highest count. **D.** Retinal flat mounts labeled for TUJ1 14-days post CONC (left). Cellpose-identified cellular boundaries were superimposed onto images using custom trained TUJ1 model (right). **F.** Cellpose accurately quantifies TUJ1+ RGCs in the CONC group but significantly undercounts TUJ1+ RGCs in the sham group (P<0.001 for both significant comparisons, n=5 eyes per group, n=3 human evaluators assessed each image, two-way ANOVA, Fisher’s LSD post-hoc). Scale bar = 20 μm.

BRN3A (POU4F1) is another commonly used RGC label^3^. BRN3A is specifically localized in RGC nuclei, unlike the cytoplasmic marker of RBPMS, and the broader label of TUJ1. To see if Cellpose could identify BRN3A+ RGCs, we developed a preliminary BRN3A model by training 8 BRN3A+ images from retinal flat mounts (Figure S4). Testing the model on separate BRN3A+ images showed Cellpose had a high degree of accuracy in identifying BRN3A+ RGCs. Therefore, Cellpose can identify RGCs using multiple labels (RBPMS, TUJ1, and BRN3A), however, this identification is most accurate when the label is specific for the RGC soma or nucleus (RBPMS and BRN3A).

### 3.4 Custom Cellpose Models and Script for RGC Characterization

Cellpose’s ability to segment RGCs based on their immunohistochemically defined boundaries opens the potential for additional analyses to be completed in tandem with RGC counts. One important analysis commonly completed to evaluate overall RGC health is measuring RGC somal size or somal area^47,55,56^. Therefore, we evaluated whether the custom script could analyze Cellpose outputs and calculate RGC soma size comparably to human analyzers. The human analyzers calculated soma size using a previously published method in ImageJ^47^ (Figure 5). Soma size calculation using Cellpose’s RGC segmentation results was included as a custom feature in the developed script. It is important to note that both methods exclude cells that cross the border of the image, meaning only cells present in their entirety are included. There was a positive correlation between the script’s calculation of soma size (running the Cellpose RBPMS model) and the average human evaluator’s calculated soma size (Figure 5A, R^2^ = 0.6283). However, this correlation was lower than anticipated. To understand why there was a difference between the Cellpose’s soma size calculation and the average human, we assessed how ImageJ and Cellpose determined cellular boundaries (Figure S5). The ImageJ process for analyzing somas involves using a watershed separation algorithm instead of segmentation. This process is inconsistent in its ability to identify RGC borders, sometimes resulting in borders being drawn that are not biologically accurate. For example, the watershed algorithm resulted in cells where excess area was included (Figure S5C), additional cells being counted that should not have been (Figure S5D), cells with too little area being counted (Figure S5D), and fragmented cell boundaries (Figure S6E). In contrast, Cellpose is an algorithm specifically designed to segment cells. In the same examples (Figure S5C-E), Cellpose identified cellular boundaries in a manner comparable to how a human would define cell boundaries in the image. Taken together, Cellpose appears capable of more closely matching the cellular boundaries defined by humans, and thus likely represents a more accurate soma size calculation when compared to prior methods using ImageJ.

**Figure 5:**
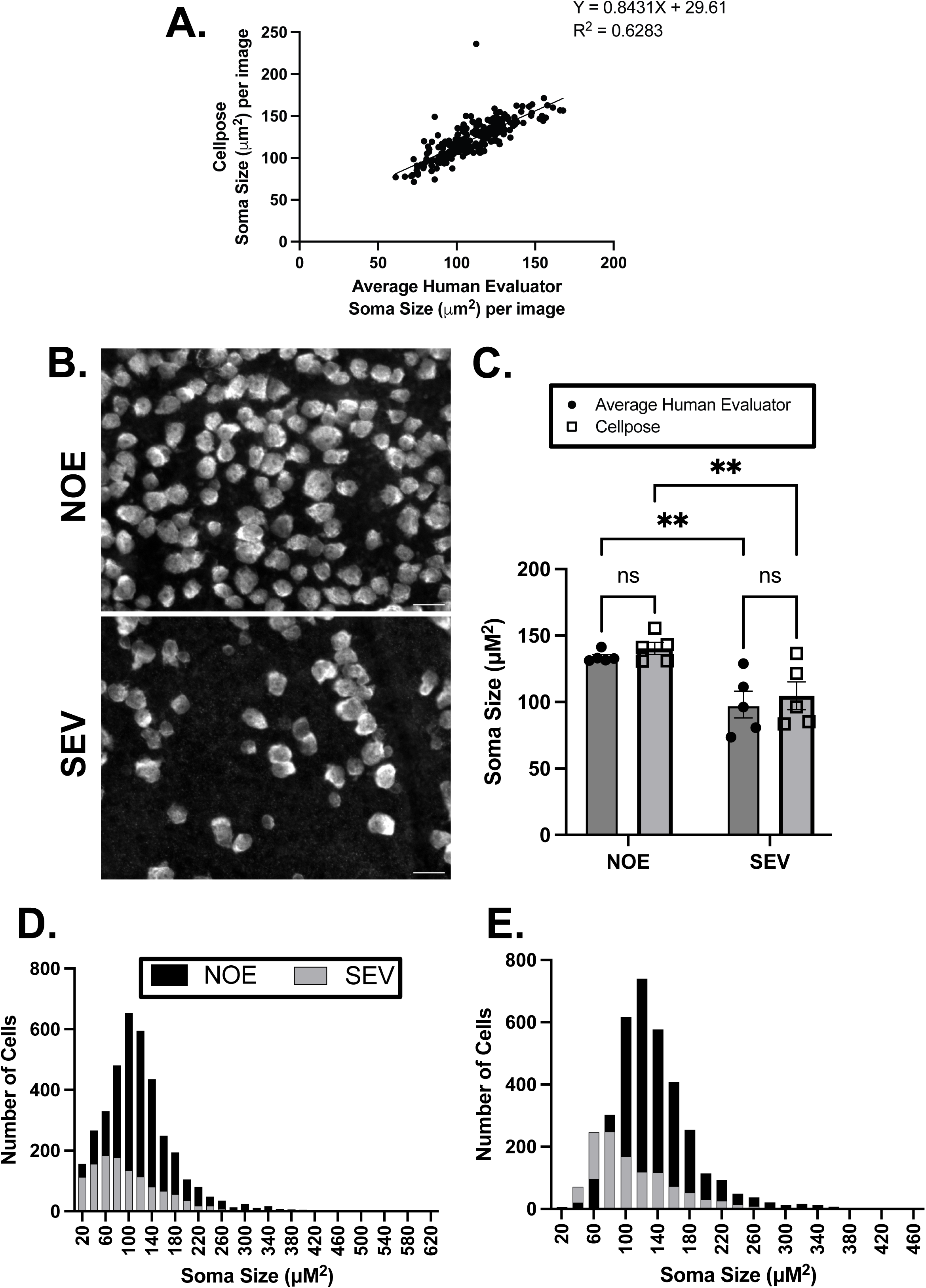
Custom Script enables automatic quantification of RGC soma size. **A.** Simple linear regression comparing the script’s quantification of RBPMS+ RGC soma size compared to the average human evaluator using ImageJ to quantify the same images. **B.** Example RBPMS labeled RGCs from 12-month DBA/2J retinal flat mounts corresponding to optic nerves with either NOE or SEV degeneration. Scale bar = 20 μm. **C.** Cellpose and ImageJ quantification of somal area (P<0.01 for both significant comparisons, n=5 eyes per group, n=2 human evaluators assessed each image, two-way ANOVA, Fisher’s LSD post-hoc). **D.** ImageJ quantification of individual RGC soma sizes across the two DBA/2J conditions. RGCs from the NOE group are shown in black and RGCs from the SEV group are shown in gray. **E.** The script’s quantification of individual RGC soma sizes across the two DBA/2J conditions. RGCs from the NOE group are shown in black and RGCs from the SEV group are shown in gray.

**Figure 6:**
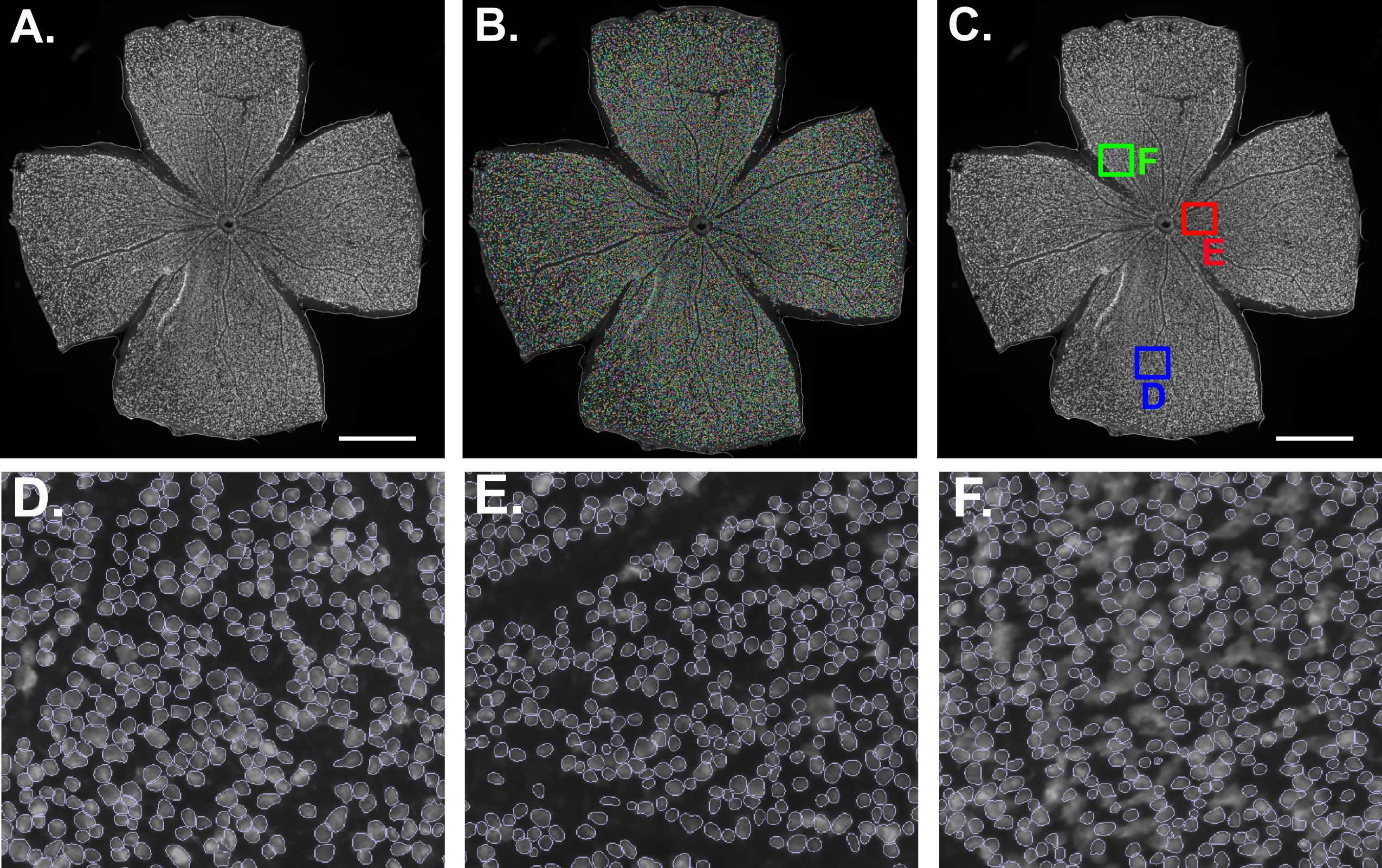
Cellpose can quantify RGCs from an entire retina. **A.** Keyence epifluorescent image of RBPMS+ RGCs in a retinal flat mount. **B.** Cellpose-identified cellular boundaries were superimposed onto the same image after application of custom-trained Cellpose model. **C.** Identification of regions on the retina for further analysis in D-F. **D, E.** A zoomed in region of the whole mount, with white lines outlining all the places the full-retina Cellpose model identified a cell. **F.** A zoomed in region of the whole mount, showing an area where the full-retina Cellpose model did not have an accurate identification of RGCs. Scale bar = 1 mm.

We then sought to assess whether the script could be used to characterize RGC soma size changes during an injury state. Prior studies have observed reduced RGC soma size in aged DBA/2J retinas^41,56^, but not in mice that have undergone CONC^55^. Therefore, we assessed whether RGC soma sizes shrunk in 12-month DBA/2J animals with corresponding SEV nerves compared to 12-month DBA/2J animals with corresponding NOE nerves (Figure 5B-E). Human raters and the script both found that eyes with a corresponding SEV nerve had RGCs with significantly smaller retinal somal area (Figure 5C). When each individual RGC was assessed, human quantifications using ImageJ showed that there was an overall shift in somal area toward smaller soma sizes in the SEV DBA/2J eyes (Figure 5D). This trend was replicated with the Cellpose data (Figure 5E). Together, this suggests that the script can characterize soma size changes in injured 12-month DBA/2J retinas.

Finally, many studies quantify the RGC population across the entire retina, instead of sampling specific regions across in retina^13,15,17,18^. This allows for analysis of changes that may be spatially isolated or differential across the entire retina. Although not the primary focus of our study, initial tests showed that quantifying an entire retina was feasible using Cellpose (Figure 6). Whole retina images were obtained using a Keyence BZ-X800 epifluorescent microscope. A preliminary Cellpose model was trained on small a region of one image containing approximately 400 cells and spanning from the peripheral edge of the retina to the center of the retina. A separate whole retina image labeled with RBPMS (Figure 6A) was used to test the performance of the preliminary model. Even with relatively modest model training, the Cellpose model was able to accurately identify the majority of RBPMS+ RGCs in the entire retina (Figure 6B). Regions in both the periphery and the central retina were able to be accurately identified (Figures 6C-E). Cellpose estimated this retina had 50,520 RGCs, which is in-line with total RGC counts in previously published reports^57,58^. However, the model was not perfect, and there were areas that were not accurately quantified (Figure 6F). Given that it is recommended that 500-1000 cells are used to train a given model to reach maximal segmentation accuracy^22^, this suggests that with additional training, Cellpose would be capable of accurately segmenting and quantifying RGCs from an entire retina.

## 4. Discussion

RGC quantification and characterization has historically been done by hand, which limits the quantity and potential quality of data collected, as hand-counts are time consuming and are prone to bias. Cutting edge software has made it possible to automate counting processes. Here, we present a custom-designed script and models using the open source, cell segmentation software Cellpose that allows users to characterize RGC somas. The software developed provides an efficient, comprehensive, all-in-one platform for RGC somal analysis. Per image, our script quantifies cell number, average luminance, average soma size, and average soma length and width (Figures 1-5). Performance of our custom trained RBPMS model is directly comparable to the results from human counters (Figure 3). Finally, our custom trained models also demonstrate that Cellpose can quantify entire retinas (Figure 6). Together, this data shows that Cellpose combined with our custom scripts and models, is a powerful software for quantifying and characterizing RGC somas in a variety of settings.

There are important features of Cellpose that make it a powerful analysis tool. First, the Cellpose algorithm can accurately segment cells. Prior methods of cellular segmentation have been capable of distinguishing cells from each other, but many use either thresholding or placement of an artificial shape or dot over identified cells^11–19^. While thresholding can provide sufficient data on many cells, it becomes problematic when cells are clumped closely together. Further, while placing a dot or shape above each identified cell allows for quantification of how many cells are in a particular image, metrics such as soma size would then have to be quantified using additional analyses. Cellpose improves upon these prior systems, accurately identifying cellular boundaries without thresholding and providing the means to not only quantify RGCs, but also to characterize their soma sizes. The addition of characterizing RGC somas is important, as shrinkage in RGC soma size has previously been linked to injury and metabolic stress^59,60^. Quantification of soma size, therefore, can provide information about RGC health that cannot be obtained by quantification of RGC number alone. Our custom script leverages this improvement to measure multiple parameters on each dataset, saving time and standardizing the analysis workflow (Figure 1).

Outside of the functionality of our custom script, it is also important to note that our custom trained models provide a high degree of accuracy-comparable to the average human analyzer. For RBPMS+ RGCs, our script (running the RBPMS Cellpose model) had a correlation coefficient of 0.9961 with the average human counter (Figure 2), a coefficient that is in line with many previous studies with an R^2^ value of 0.95 or greater^11,13–15,18,19^. Further, our model was able to accurately identify RGCs with many different injury statuses, including in aged DBA/2J retinas and in CONC retinas (Figure 3). Cellpose was less accurate in quantifying TUJ1+ RGCs (Figure 4). This result is not particularly surprising, as other studies have also failed to obtain accurate quantification of TUJ1+ RGCs without additional image processing^12^. Prior studies have noted that TUJ1+ immunolabeling of retinas can be inconsistent within and among samples, which was the case in our dataset^12^. Additionally, TUJ1 labels the processes of RGCs, which can make identifying somas accurately a challenge. Together, the inconsistent labeling combined with inclusion of processes likely contributed to Cellpose’s inaccurate quantification of TUJ1+ RGCs in healthy retinas, where many RGCs are present and where RGC processes often overlap with other nearby cells. Injured retinas commonly have fewer processes and more space between RGC somas, which makes their identification easier and could explain why our model was able to more accurately quantify injured retinas compared to healthy retinas.

Finally, our script allows for multiple supplemental analyses to be completed, such as quantifying features of soma size or performing quantifications across an entire retina. Consistent with our prior work, Cellpose and human-driven ImageJ analysis both showed that RGC soma size is reduced in 12-month DBA/2J retinas corresponding to nerves with SEV degeneration (Figure 5)^41^. This finding is in line with many prior studies suggesting ocular hypertensive insults lead to a shrinking or selective loss of larger RGCs^61–64^. Quantifying RGCs across the entire retina is also important to the study of RGC somas. The RGC death seen with CONC is extensive and seen throughout the retina. In contrast, RGC death in ocular hypertensive DBA/2J retinas can occur in a radiating wedge-shape from the optic nerve^65^, thereby affecting areas of the retina differently. Other models, such as some variations of the microbead model of experimental glaucoma^66,67^ or injections of endothelin-1^33,34,68,69^ or TNF-α^70,71^, cause milder degrees of RGC death, for which imaging the whole retina could provide a powerful platform to better gauge the changes occurring to RGC somas. Additionally, the large quantities of RGCs present within a retina make it impractical to count and characterize RGC somas manually. The work we present here demonstrates that quantification and characterization of RGC somas is possible within the Cellpose platform using our custom script (Figure 6).

In conclusion, our custom designed script and models use Cellpose to provide an all-in-one analysis platform for RGC somas. The models have a high degree of accuracy, comparable to human analyzers, and the script allows for multiple analyses to be run simultaneously, saving investigators time. Cell quantification and characterization are critical outcome measures for the study of optic neuropathies, and many other fields. However, this process can be exhaustive, tedious, and prone to bias. The application of Cellpose, as we have presented here, improves the reproducibility and efficiency of research investigating RGC somas.

## Acknowledgements

The authors would like to acknowledge the Center for Advanced Light Microscopy and Nanoscopy at the University of Rochester and Dr. V. Kaye Thomas for their excellent technical support. Further, the authors thank Dr. Juliette McGregor for use of her computational resources.

## Data Availability

The script, models, and all associated information is publicly available at https://github.com/abigailbishop/LibbyLabCellpose.

---

**Figure S1:**
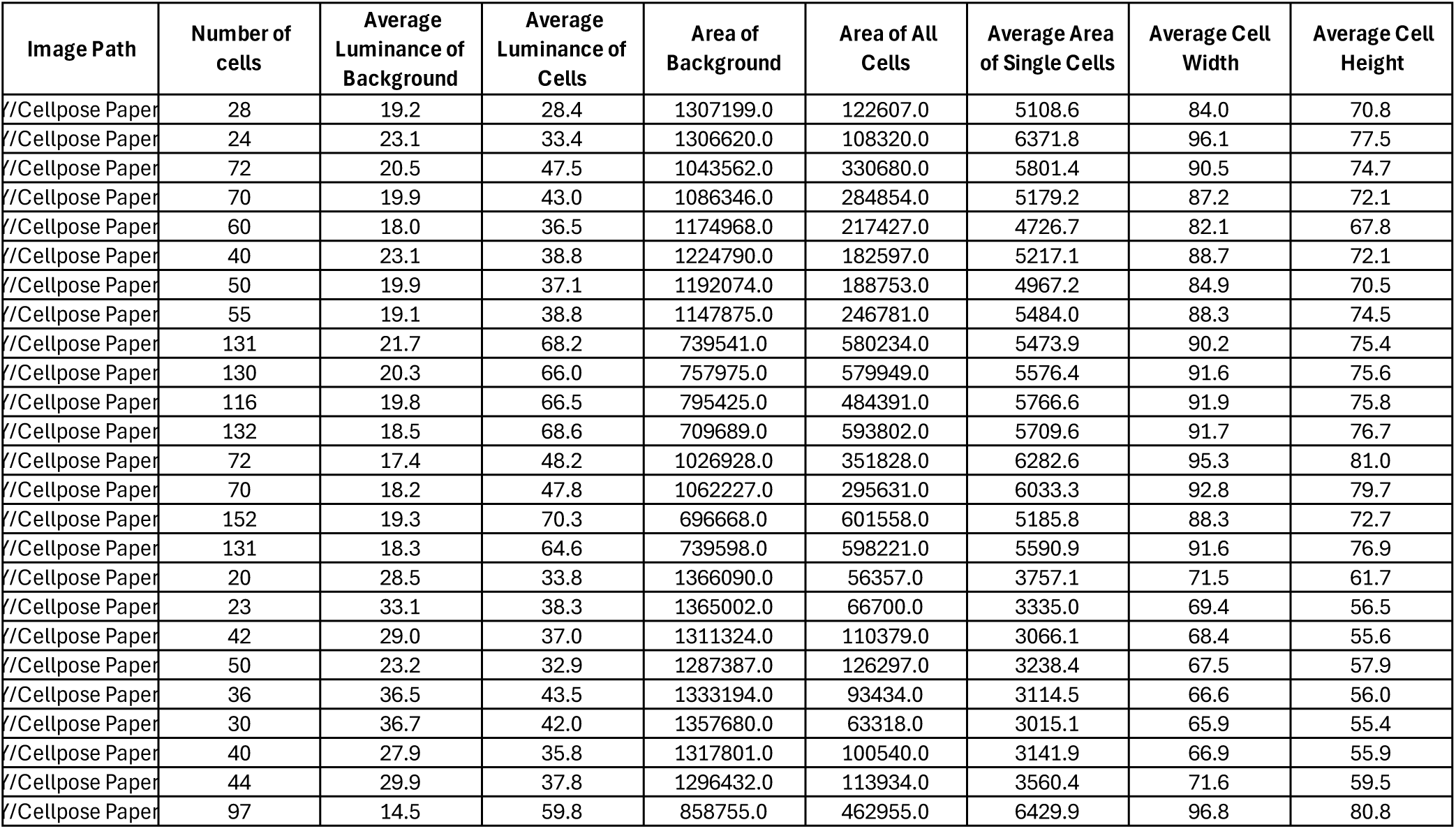
A sample .CSV file output from Cellpose. In the first row, the path to the model used to analyze the images is given. In the second row, the cell diameter (in pixels) assumed by the model is given. In the third row, the output metrics from the script are listed. Each additional row represents the output for an individual image. Metrics such as area, width, and height are all represented in pixels, though an optional argument can be used to provide a conversion factor and automatically convert the data to a desired unit when running the script.

**Figure S2:**
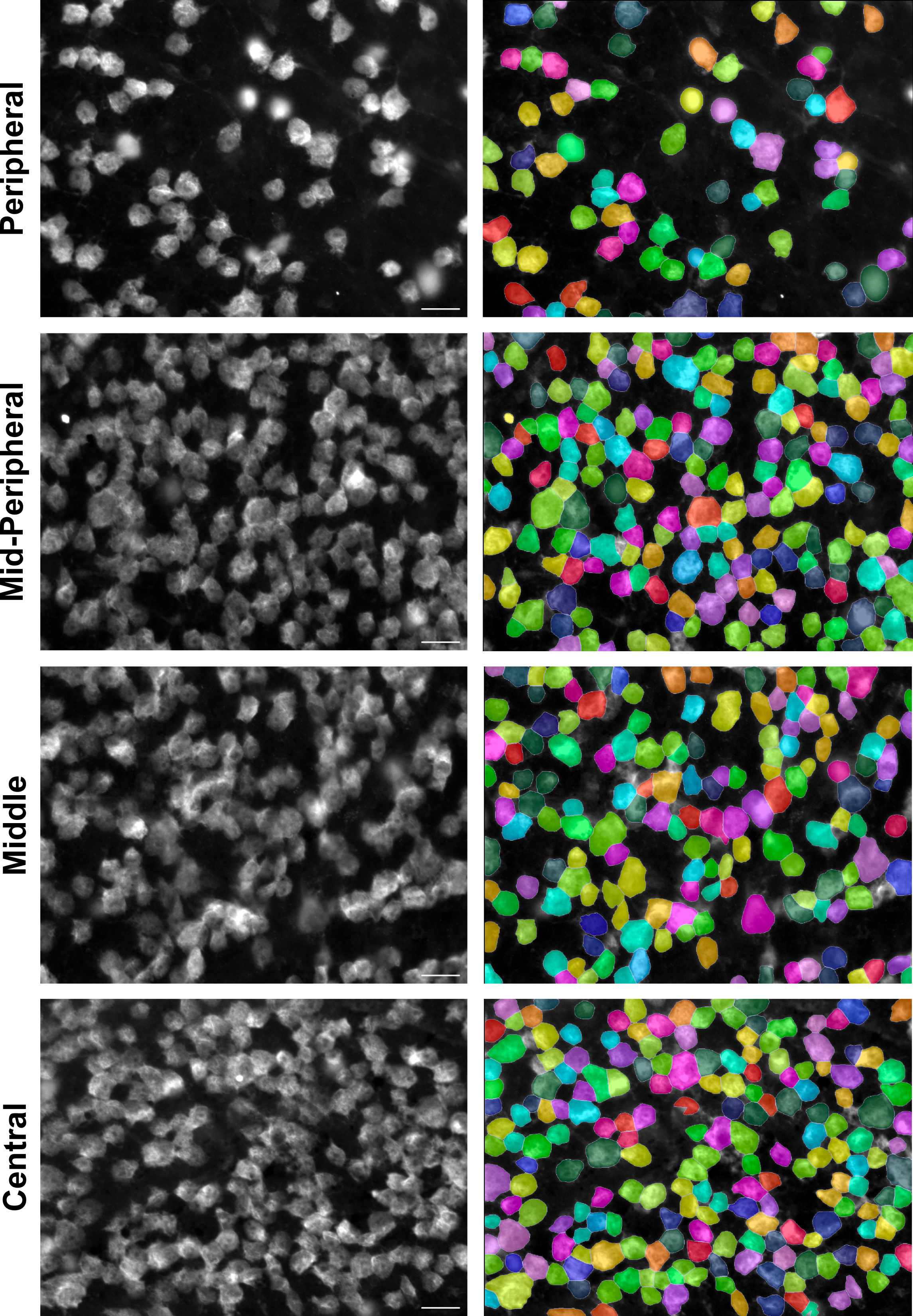
Custom RBPMS Cellpose model can distinguish RGCs at various locations in the retina. Flat-mounted retinas were labeled for RBPMS, and images were acquired at 40X magnification at various regions throughout the retina. Example RBPMS-labeled RGCs in each region shown in their original image (left) and with superimposed Cellpose-identified cellular boundaries (right) after running the custom-trained RBPMS Cellpose model. Scale bar = 20 μm.

**Figure S3:**
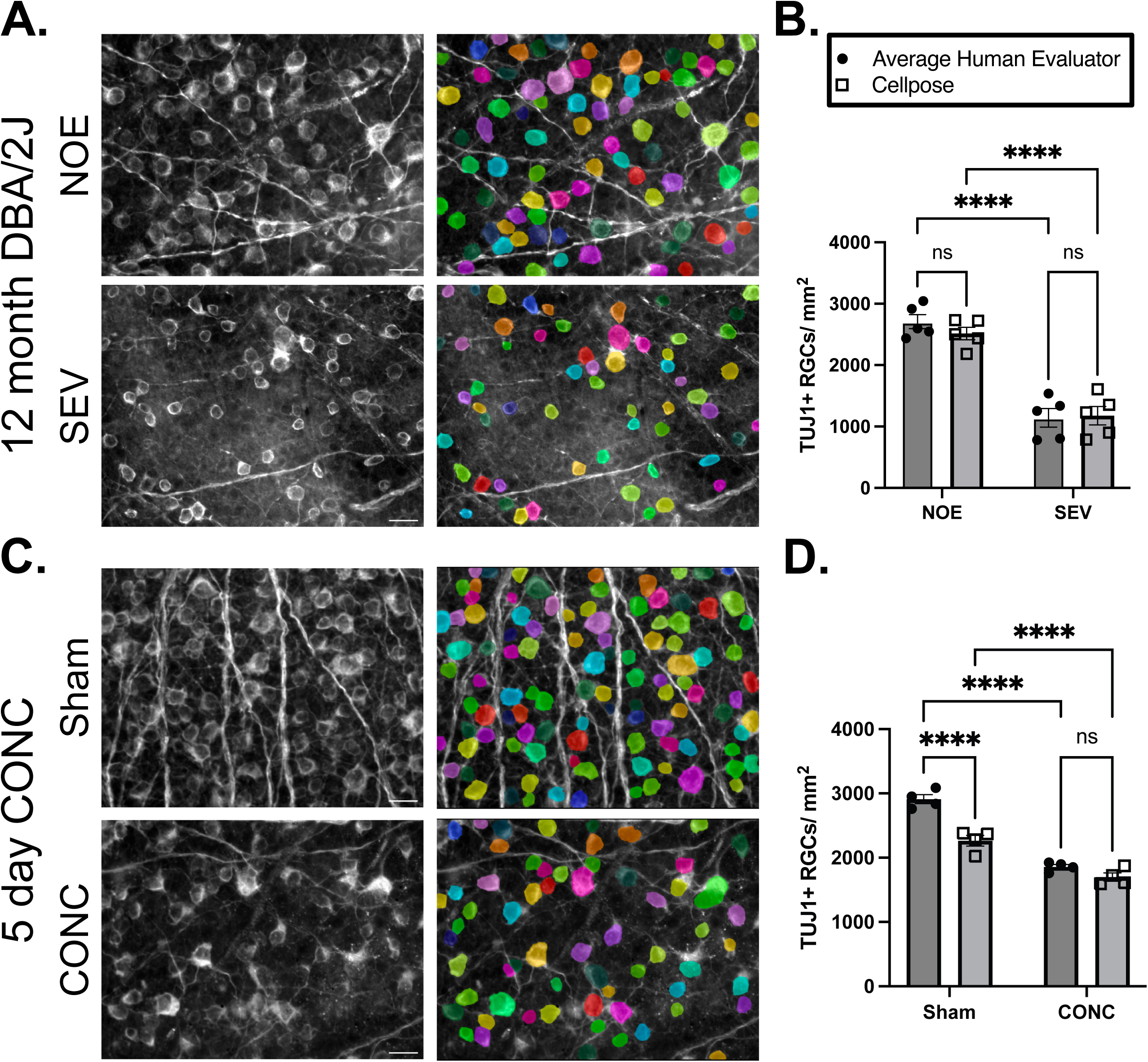
Custom TUJ1 Cellpose model’s performance with 12-month DBA/2J and 5-day CONC data. **A.** Retinal flat mounts from eyes corresponding to optic nerves with NOE or SEV degeneration from 12-month DBA/2J animals. Flat mounts were labeled for TUJ1 (left). Cellpose-identified cellular boundaries were superimposed onto images after quantification using custom trained TUJ1 model (right). **B.** Cellpose quantifies RGC density in NOE and SEV groups in a manner comparable to human evaluators (P<0.001 for both significant comparisons, n=5 eyes per group, n=3 human evaluators assessed each image, two-way ANOVA, Fisher’s LSD post-hoc). **C.** Retinal flat mounts from eyes receiving CONC or sham injury, 5 days post-procedure. Flat mounts were labeled for TUJ1 (left). Cellpose-identified cellular boundaries were superimposed onto images after quantification using custom trained TUJ1 model (right). **D.** Cellpose accurately quantifies TUJ1+ RGCs in the CONC group but significantly undercounts TUJ1+ RGCs in the sham group (P<0.001 for significant comparisons, n=4 eyes per group, n=3 human evaluators assessed each image, two-way ANOVA, Fisher’s LSD post-hoc). Scale bar = 20 μm.

**Figure S4:**
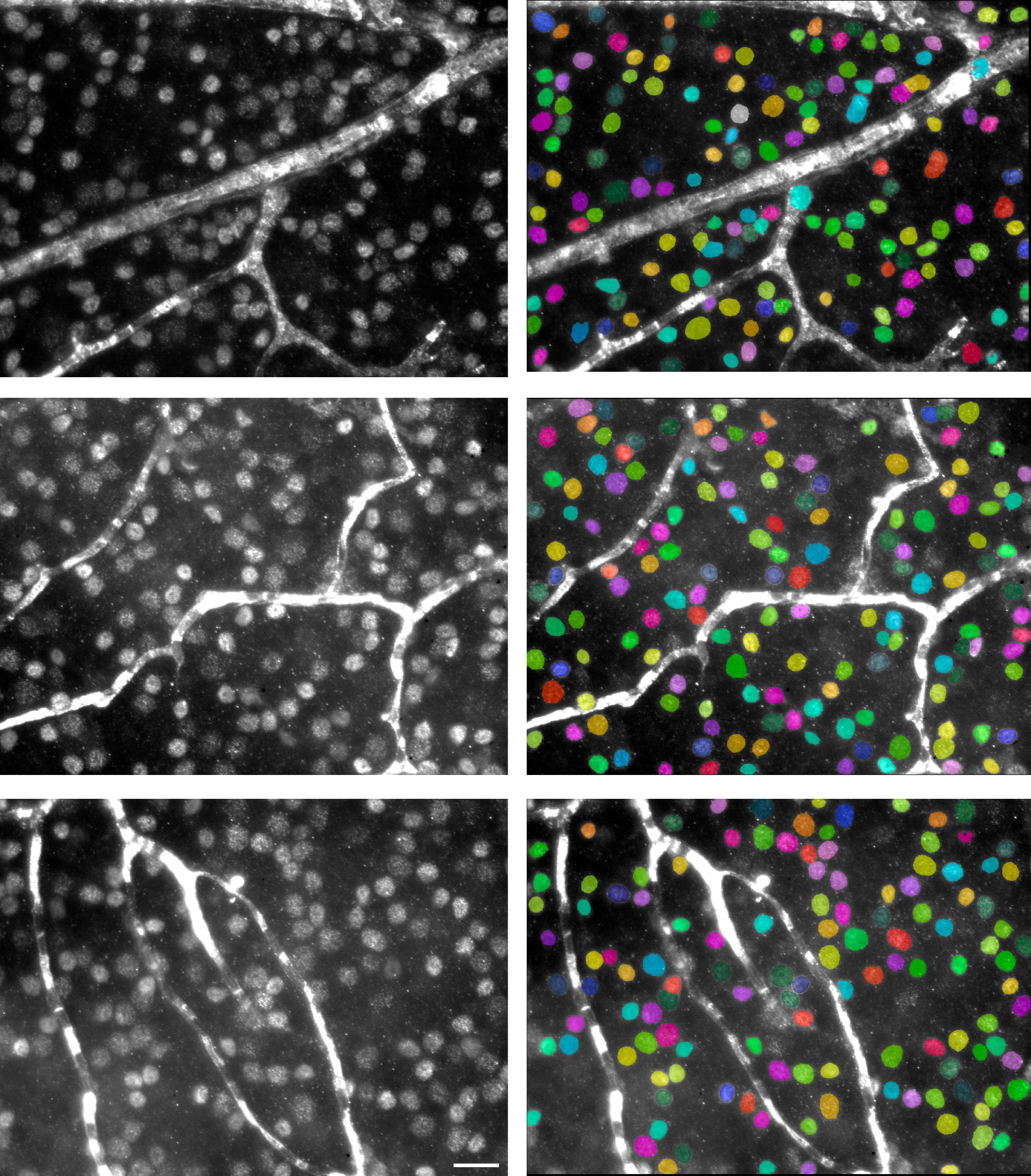
Custom BRN3A Cellpose model accurately identifies BRN3A+ RGCs. Flat-mounted retinas were labeled for BRN3A, and images were acquired at 40X magnification. Example BRN3A-labeled RGCs from three different retinas are shown in their original image (left) and with superimposed Cellpose-identified cellular boundaries (right) after running the custom-trained BRN3A Cellpose model using the script. Scale bar = 20 μm.

**Figure S5:**
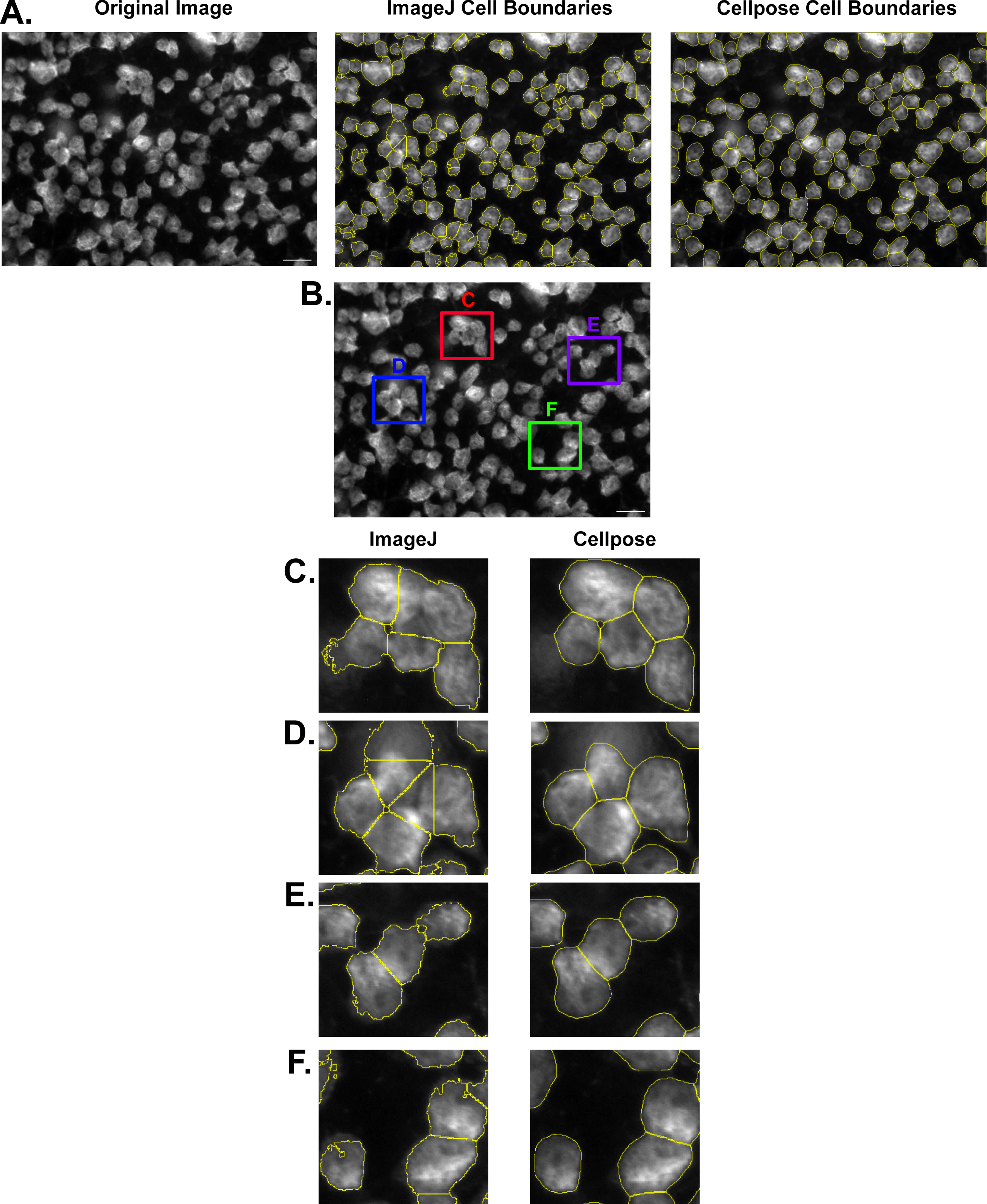
Comparison of ImageJ and Cellpose Soma Size Analyses. **A.** *Left:* Retinal flat mount with RGCs labeled for RBPMS. This image is used throughout the rest of the figure. *Middle:* The same retinal flat mount with yellow lines indicating cellular boundaries as identified by the ImageJ soma size analysis pipeline. *Right:* The same retinal flat mount with yellow lines indicating cellular boundaries identified with the Cellpose analysis pipeline. **B.** Regions of the retinal flat mount that are highlighted in figures C-F. **C-F.** Examples of regions of the retinal flat mount comparing the boundaries identified by ImageJ and Cellpose, demonstrating that Cellpose identifies boundaries between cells consistent with human-identified boundaries compared to ImageJ.

